# Palaeoproteomic deconvolution of physical and genetic collagen mixtures

**DOI:** 10.64898/2026.06.17.732552

**Authors:** Ian Engels, Tristan Dedrie, Synnøve M. Saugen, Simon Van de Vyver, Thijs R. A. Vandenbroucke, Kevin Di Modíca, Jan Decher, Alice Toso, Dieter Deforce, Simon Daled, Alexandra Burnett, Grégory Abrams, Maarten Dhaenens

## Abstract

Species identification in palaeoproteomics relies on genome-derived protein sequences which are often poor-quality, and lacks tools to cope with multi-species samples. Here, we address both challenges through the analysis of ‘physical and genetic mixtures’. Species that are absent from our database are considered a ‘genetic mixture’, i.e. a patchwork of peptides from closely related species. Inversely, various overlapping peptide stretches allow us to resolve complex ‘physical mixtures’. This is benchmarked by analysing physical mixtures of modern bone fragments, including genetic mixtures. We illustrate the impact of our approach via a rapid and high-throughput analysis of >2500 bone fragments, revealing the Eemian-era faunal environment around Scladina Cave, including the first *Palaeoloxodon antiquus* identified at this site.

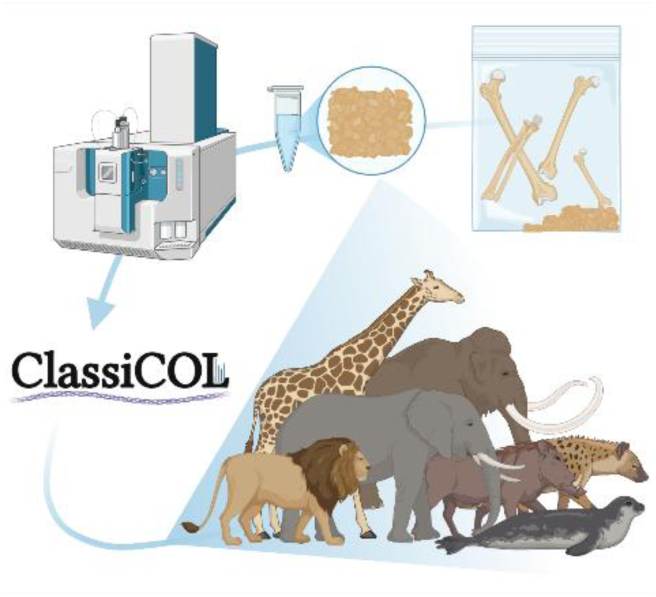

## Introduction

Identifying well-contextualized vertebrate remains from archaeological and palaeontological sites is essential for reconstruction of ancient ecosystems, and human-fauna interactions (*1*, *2*). However, the vertebrate record available to researchers is strongly shaped by differential taphonomy (*3*, *4*) and analytical constraints from highly fragmented or morphologically ambiguous bones that are frequently unidentifiable using osteomorphological characteristics. These poorly diagnostic specimens often outnumber identifiable remains and closely related or previously unrecorded taxa may therefore remain undetected or misidentified (*5*). As a result, substantial portions of vertebrate assemblages, often curated as bulk, indeterminate material, constitute an underexplored archive of biodiversity that could refine interpretations of past ecosystems and ecological change (*6*, *7*).

Within the field of palaeoproteomics, a very fast, relatively cheap and effective analytical technique is commonly used to analyse large collections of bone fragments: ZooMS (*8*). Herein, each individual bone fragment is sampled, its proteins digested with trypsin and analysed using MALDI-MS. Due to their structural function, collagens are well-embedded in the mineralized bone matrix, protecting them from degradation over time and making collagen the most prominent biomolecule within ancient bones (*9*). Collagen is the target of ZooMS analyses, enabling species to be identified based on taxon-specific protein masses. Unfortunately, ZooMS is inherently incapable of disentangling mixtures (e.g. contaminated bones) or identifying new species that are not represented in marker databases. Such discriminating capabilities would exponentially increase the throughput and impact of palaeoproteomics.

Naturally disintegrated bone dust, collected from plastic bags in which unidentified bones are stored in bulk, provides a single sample that captures the whole species content of the bag. In this way, whole collections could be screened to find those bags that contain potentially interesting fragments of e.g. rare or extinct animals, including elusive Hominin remains. Likewise, coprolites (*10*, *11*) can provide a snapshot of multiple animals consumed by a predator or scavenger within a very narrow window of time. This not only tells us about the presence of certain animals in the area but also the interactions between prey and carnivore (*12*). Additionally, bone tools or lithic tools (*13*), or artefacts to which animal-based glues were curatorially applied, could still yield collagens of numerous species. Identifying organic surface alterations is important as artefacts can be selected for carbon dating only when animal glue is absent. Finally, aDNA and proteins have both been found in soil (*14*, *15*), implying that the capability to proteomically identify mixtures could allow palaeoenvironmental reconstruction of entire caves using soil samples from all relevant layers. Many of these applications are minimally invasive, making them especially valuable in the contexts of archaeology and palaeontology.

Liquid Chromatography-tandem Mass Spectrometry (LC-MS/MS) has emerged as a powerful and information-rich acquisition strategy in palaeoproteomics (*16–19*). LC-MS/MS was developed to annotate proteins in conventional proteomic experiments, where a measured set of peptide (digested protein) masses need to be assigned to the proteins from which they originated, while considering several similar homologues and paralogues in the database. A complication inherent to this mathematical challenge in proteomics analysis is ‘*the protein inference problem’* (*20*, *21*). In palaeoproteomics, the challenge is orders of magnitude larger and is defined as the ‘*species inference problem’*: finding the most likely protein sequence, not from a few homologue proteins within one species but of hundreds of very similar orthologues from different species. The species inference problem has only recently been approached more generally (via UniPept) (*22*) and more recently tackled in respect to collagen (via ClassiCOL) by the current authors (*11*).

In its first iteration, ClassiCOL was not yet capable of complex multi-species differentiation. We propose to approach species classification from a novel perspective, with the following distinctions: a **physical mixture** is a sample composed of multiple species, while a **genetic mixture** is a sample from a single species not present in the database, but which can be considered a mixture of closely related species. We reason that the first is reflected in the data by collagen sequence stretches that are persistently covered by several peptides derived from different species. Inversely, the latter is reflected in the data as a collagen sequence that can only be patched together by non-overlapping collagen peptides from different (closely related) species that are present in the database. Many extant mammalian species lack or have truncated protein sequences and for most no experimental (mass spectrometry) evidence is available to validate the predicted sequences. There is additionally a clear database bias towards species from the Northern Hemisphere, while taxonomic families from the Southern Hemisphere are often poorly represented. Many extinct animals are also absent from protein databases. This new form of theoretical sequence reconstruction is therefore widely applicable.

We have embedded this novel perspective in ClassiCOL_v2, which can now approximate the species present in complex mixtures, consisting of extant and extinct species, in varying abundances and states of preservation. We also correct prediction errors and gaps in the CollagenDB without the need for *de novo* sequencing approaches, demonstrating these capabilities by reconstructing the faunal environment of Scladina Cave, Belgium, based on bone dust taken from 24 bags containing >2500 bone fragments.

## Results

### Collagen for taxonomic classification

In our previous report (*11*), we described a species inference tool to classify species based on LC-MS/MS data searches with the CollagenDB. The first iteration of ClassiCOL was capable of limited mixture analysis, producing sunburst plots displaying several high-scoring potential candidate taxa. This enabled the identification of both Bovinae and *Ursus* in a >40,000-year-old cave hyena (*Crocuta crocuta spelaea*) coprolite. However, the interpretation of the sunbursts was complex and prone to user error, especially when interpreting mixtures. Therefore, we developed a dedicated algorithm to resolve complex physical and genetic mixtures. These analyses are performed at an average 10x reduction in computational time and memory usage when compared with ClassiCOL_v1 (**Figure S1**).

We first investigated the theoretical power of using only two proteins - COL1α1 and COL1α2 - to resolve phylogenetic relatedness over varied timescales by constructing a phylogenetic tree *in silico* directly on the collagen sequences for the species in the CollagenDB (*11*). **Figure 1A** shows this for species belonging to the superorder Laurasiatheria. Indeed, this tree closely resembles the NCBI taxonomic tree down to the species level, showing the potential of the curated sequences in the CollagenDB to achieve very accurate classifications.

**Figure 1.**
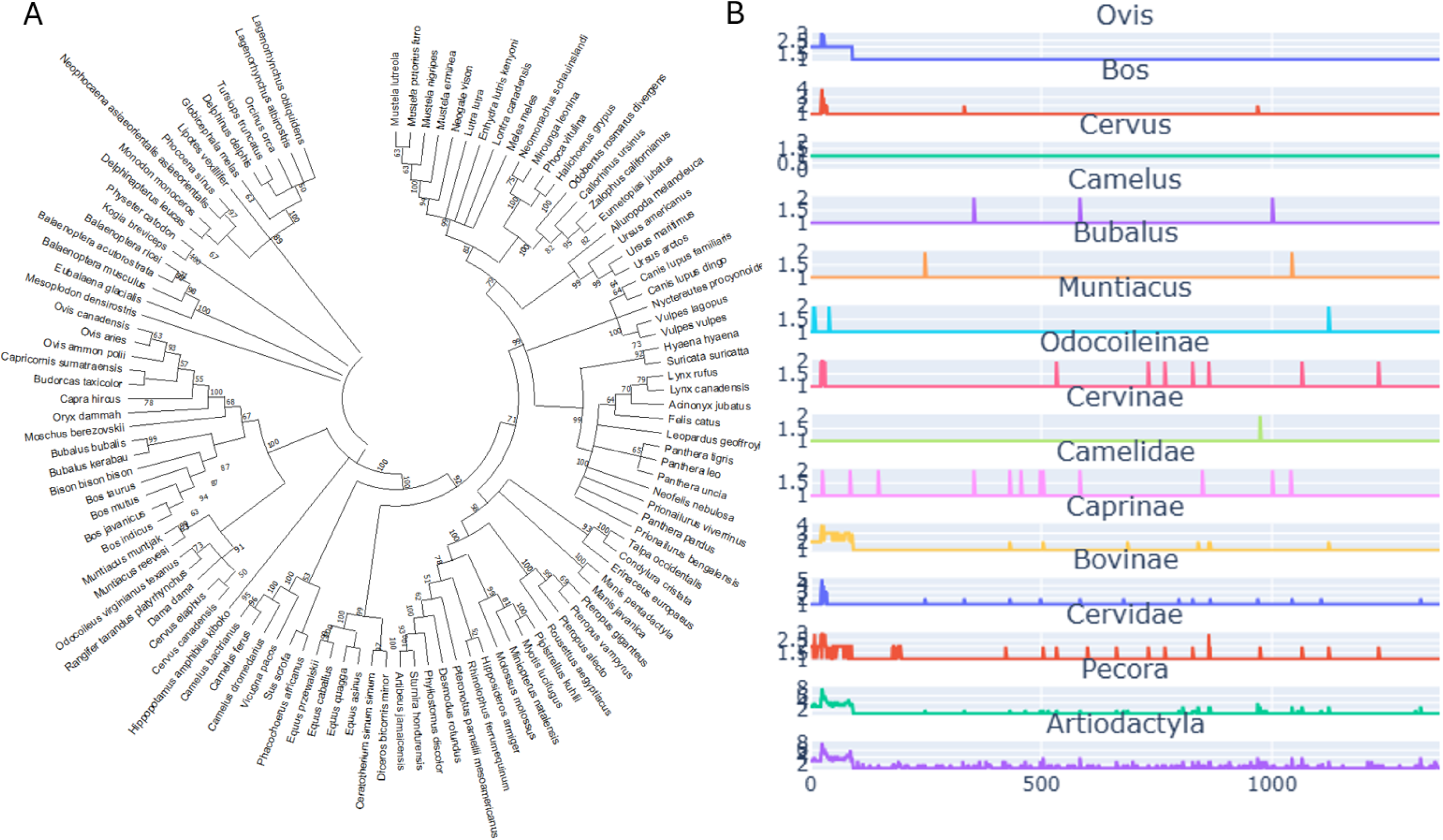
Framework for multiple-species inference using protein phylogeny and mutational space mapping. **A. The collagen phylogenetic tree resembles the NCBI taxonomic tree.** Using concatenated collagen sequences of COL1α1 appended to COL1α2, a phylogenetic tree was constructed using MEGA11 software (*23*). Species names of taxa present in the CollagenDB are shown. Distances were calculated using the Poisson method and clustered via the Neighbour-joining method. **B. The collagen mutational search space increases in association with distance from the last common ancestor.** Line plots show mutational residue locations in COL1α2 for species in the CollagenDB under their last common ancestor. X-axes show residue location at the collagen consensus sequence for the Artiodactyla taxonomic level.

Through multiple alignment of collagen sequences, the CollagenDB mutational space can be mapped onto consensus sequences at each taxonomic level, where mutational residues are denoted by X’s at higher taxonomic levels. However, when considering the entire CollagenDB, this results in long stretches of ‘X’s at high taxonomic levels, rendering the sequence unusable downstream. To avoid this, we build the mutational space on every measured sample, using the ClassiCOL output as the starting point for building a sample-specific mutational space. **Figure 1B** visualizes this approach for a sample that has Artiodactyla as its LCA.

The multiple sequence alignment can be used to define the uniqueness of detected amino acid residues for extant relatives present in the database and therefore the probable ancestral origin of the sample. Each of the detected potential mutational residues are therefore scored for *recentness* of the mutation according to their ancestral lineage and *uniqueness* compared to other candidates in a one-vs-one comparative approach. As expected, the number of mutational residues compared to any given species gradually increases towards the last common vertebrate ancestor (LCVAs) as this is the *in silico* equivalent of going back in time. With the substantial differences in generational turnover between e.g. Artiodactyla and Rodentia, their mutation rates of collagen therefore also vary considerably (**Figure S2**). By normalizing these rates separately, this approach co-ordinately accounts for less obvious processes that lead to altered evolutionary pressure on the protein.

To reconstruct genetic mixtures, we can leverage evolutionary logic to perform species inference using only identified peptides. We hypothesize that only few animo acid mutations will not change the structural function of the collagen protein and can therefore be passed on throughout their ancestry as functional proteins, i.e. they are conserved among descendant species (**Fig S2**). Once a mutation becomes stable within a lineage at a certain point in time, the chances of the same residue mutating again over a short time span is rather unlikely (*24*). Therefore, we can almost completely represent species that are unaccounted for in our database as a patchwork of related collagen peptide sequences, with few novel or unique point mutations in any given species. Note that errors in the sequence database can equally be resolved by this *peptide-patch* approach. In other words, we approach the species absent from the database as genetic mixtures of known peptides, omitting the need to include *de novo* sequencing to detect them.

### Deconvolution of genetic and physical mixtures

Building on this evolutionary logic, we attribute gaps in collagen sequence coverage to the absence of the target species in the database, rather than to unmeasured peptides. Measured peptides of closely related species can be used to ‘patch’ these gaps to create a theoretical sequence, i.e. a sequential combination of existing but not co-occurring collagen peptides, which approximates the true species more closely. By iteratively comparing each new theoretical sequence with the remaining sequences from the previous iteration, the algorithm moves upstream in the taxonomic tree towards the LCVA, in a one-vs-one approach, which shifts into a many-vs-many approach near convergence, i.e. when no more combinations are possible. This convergence ends with either an exact match to a species in the database or with a theoretical sequence which can be localized to a phylogenetic location adjacent to close relatives through sequence alignment, providing a **theoretical candidate**. When dealing with true physical mixtures on the other hand, i.e. a single sample containing multiple species, multiple explanations are found in the data for a given location in the collagen sequence. Both physical and genetic mixtures can occur simultaneously when dealing with a multiple-species sample in which at least one taxon is not in the database.

The classification made by ClassiCOL_v2 on a reference *Hippotragus niger* (sable antelope), which is not present in the CollagenDB, is shown in **Figure 2**. To allow deeper curation of the result, the algorithm generates a **taxonomic slope plot** per deconvoluted (theoretical) candidate in relation to the rest of the tree (**Figure 2C**). A steeper slope implies a smaller taxonomic distance to the next taxon. Note that to make an informed decision on the sample content, prior knowledge on the sample remains crucial. For example, in the dust from bags containing a large number of bones, setting a score threshold for selecting species candidates in the sample makes little sense, because low-scoring candidates could still provide a lead to an animal present in the sample. Conversely, it is reasonable to use a score drop-off threshold when a single bone was drilled and/or a single genetic mixture is expected. However, it is important to consider that a single bone can still be contaminated with dust or animal glue.

**Figure 2:**
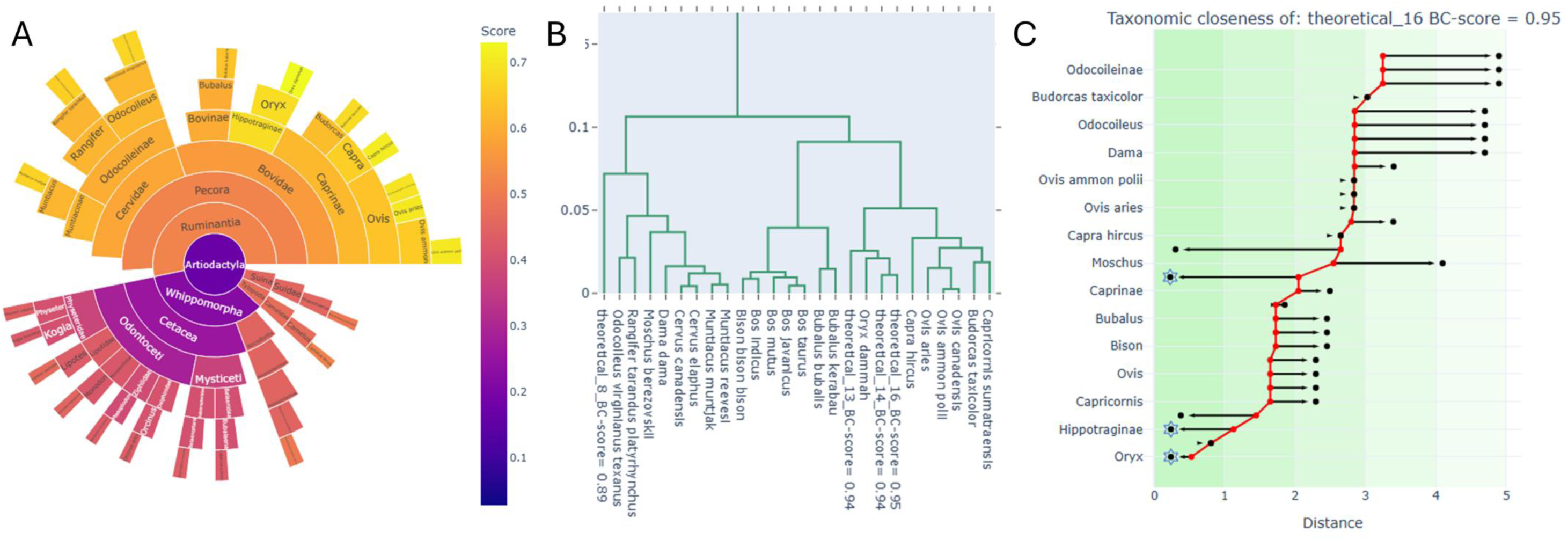
ClassiCOL output of taxonomic placement. **(A)** The sunburst plot shows the approximation of species candidates in a reference sample of *Hippotragus niger.* **(B)** Based on the LC-MS/MS measured data, a tree plot is constructed showing the taxonomic location of the theoretical species in relation to closely related taxa. ‘Theoretical_’ represents collagen sequences constructed using the peptide-patch approach on measured data from close relatives. **(C) Taxonomic Slope Plot.** The red line is a linear depiction of the distances shown in (B). It depicts the increase in ranked measured taxonomic distances from a given annotated/theoretical collagen sequence. At each stage, a horizontal arrow is added, indication the location of a species from a missing taxon, to contrast it with the distance calculated on the measured collagen peptides, i.e. the red line. An arrow to the left therefore means that the taxon most likely belongs to the main lineage, while an arrow to the right means that the taxon potentially belongs to a closely related lineage. The most likely taxonomic lineage, given the measured data, is highlighted by a blue star. By hovering over the nodes in the interactive HTML, a pop-up list of missing sub-taxa is shown (**Data S1**).

### Benchmarking genetic mixtures

There is a known bias in protein databases against representation of several family-level taxa, many being African mammals (e.g. Antilopinae, which contains *Saiga tatarica,* the saiga antelope, that was present during the Late Pleistocene in North-Western Europe (*25*)). We used representatives of missing families to benchmark the performance of ClassiCOL_v2 on genetic mixtures. First, we analysed 12 species present in CollagenDB and then 27 species that were missing from CollagenDB (genetic mixtures), representing a variety of taxonomic groups and relatedness distances. **Figure 3** demonstrates highly accurate clustering of the analysed species, whereby the red color-coding shows regions that are not present in CollagenDB, thus being considered ‘genetic mixtures’. Briefly, potential species candidates were determined by Mascot searches against the CollagenDB, followed by the ClassiCOL pipeline (including isoBLAST (*11*)). This was used as input to construct theoretical collagen sequences by clustering measured peptides, before building the taxonomic tree on measured taxonomic distances, approximating the most probable taxonomic location of the theoretical sequences.

**Figure 3:**
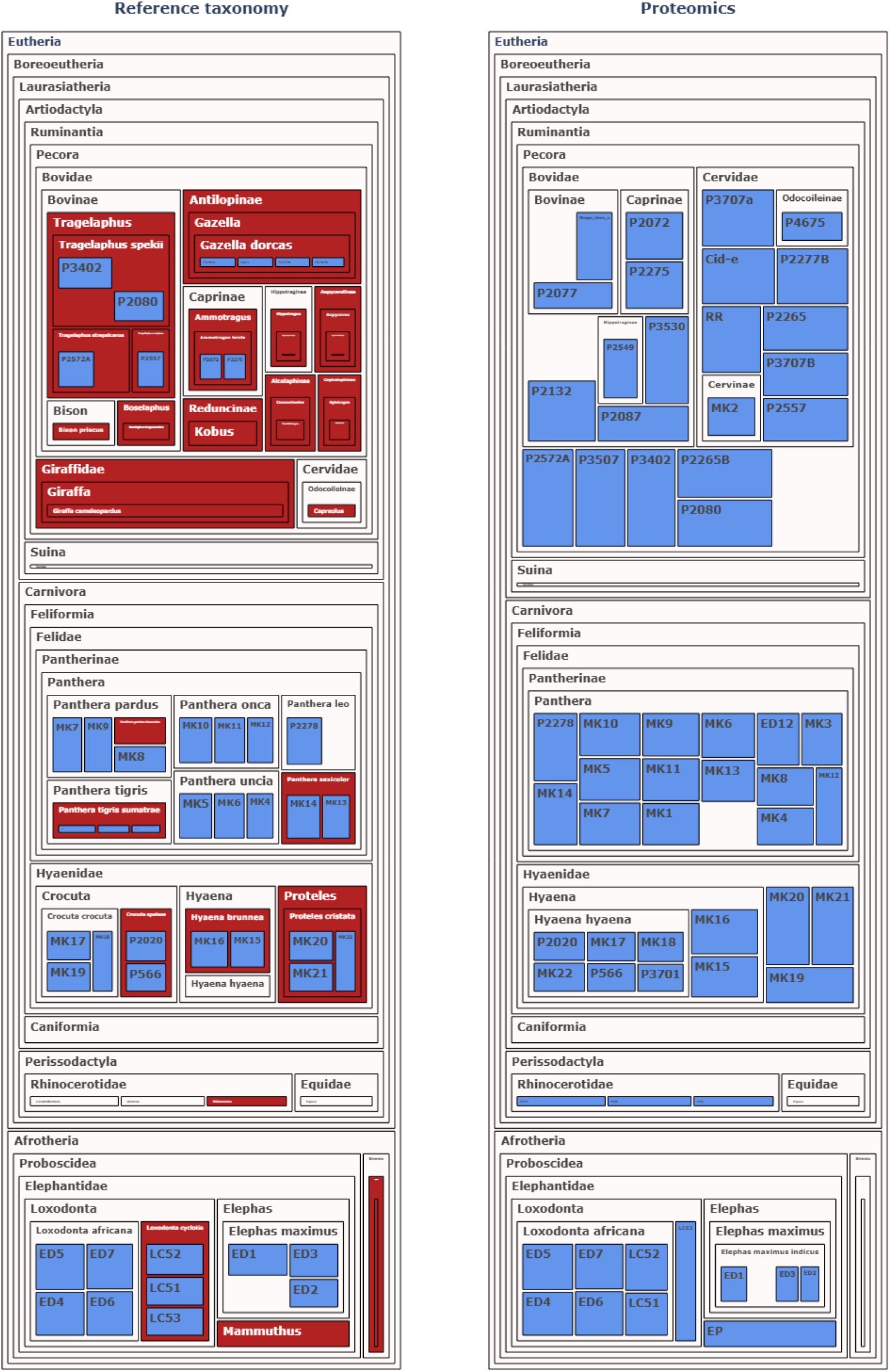
Taxonomic placement of ‘genetic mixture’ reference samples. The reference species from Museum Koenig Bonn and the UGent Sterre collection are visualised in blue in a tree map of their NCBI taxonomy classification. Taxonomic regions that are not covered in our CollagenDB are color-coded in red. Note that some samples are no longer visible because of the much greater taxonomic detail attainable in theory than by our available CollagenDB taxonomic tree shown on the right. The measured palaeoproteomics outcome of these samples is plotted in blue on the available taxonomic tree from CollagenDB, showing the attainable taxonomic location of all the samples, including the genetic mixtures. The interactive HTMLs (**Data S2**) and all tree and taxonomic slope plots are deposited in PRIDE (PXD079630).

### Benchmarking physical mixtures

To demonstrate the performance of ClassiCOL_v2 on multi-species mixtures, 14 experimental mixtures were made from bone powder drilled from the cortical surface of the left femur of ten modern specimens, spanning varying taxonomic distances, from the Ghent University Ledeganck reference collection, which were first individually analysed and matched to their osteomorphological identification **(Figure 4)**. This showed that, given the current database, *Martes foina* (beech marten) and *Mustela putorius* (European polecat) return similarly, thus making them indistinguishable from one another when both species are present in a mixture. In general, the correctly classified species in all mixtures and individual samples are ranked as the top resulting candidates, followed by lower-scoring false positives per order. Also, an absence of *Meles meles* (Badger) is noticed when it is mixed together with other species within the family Mustelidae. Similarly, *Rangifer tarandus* (reindeer) seems to be missing after deconvolution when multiple close relatives are present in the mixture.

**Figure 4:**
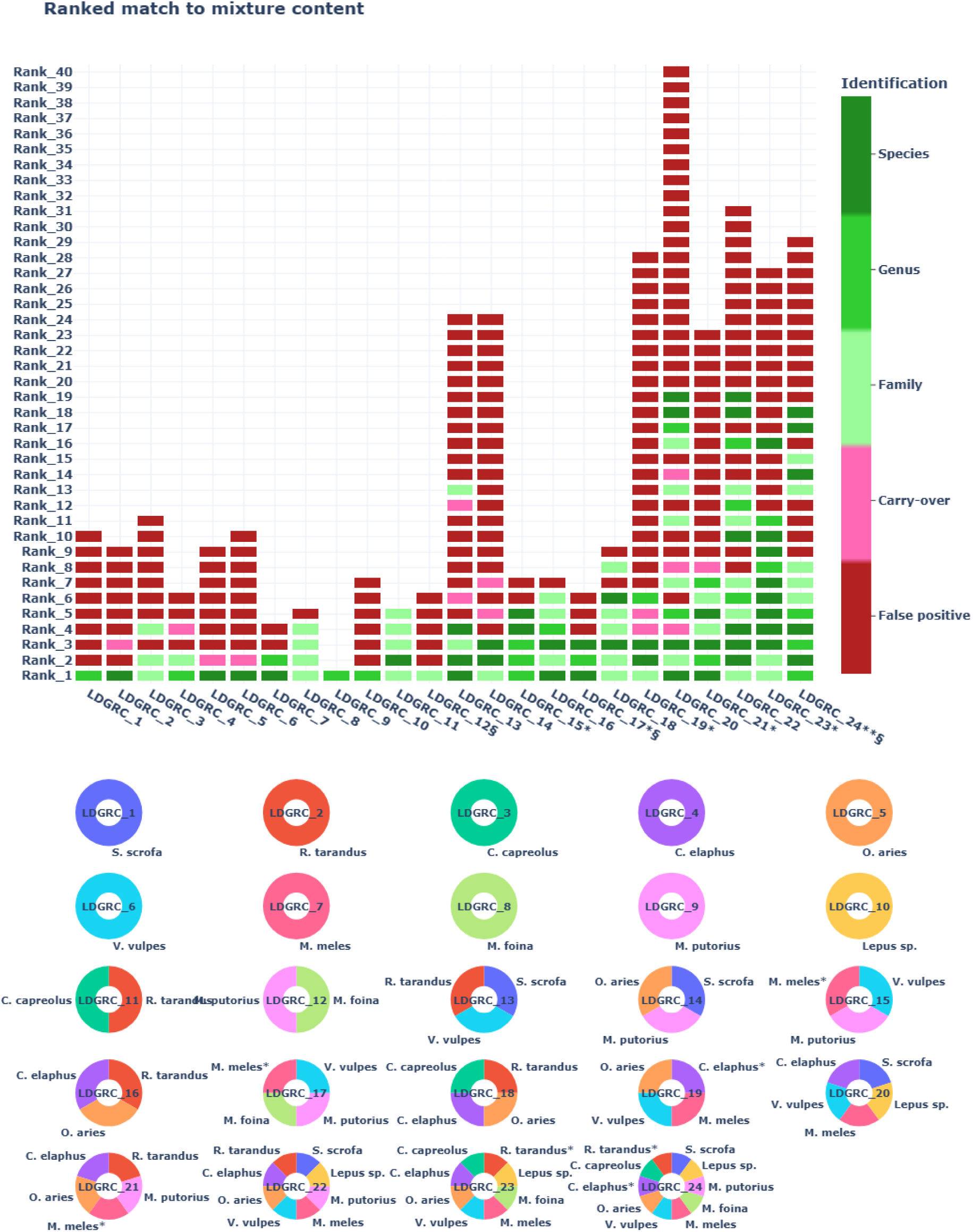
Experimental physical mixture deconvolution matches species composition. (**Top**) Ranked classification overview per sample. Taxa matching the sample content are shown in green, with dark green for species matches, towards light green for family taxon level matches. Potential instrumental carry-over species (present in the sample n-1 in the sample list) are visualized in pink, and false positive or ambiguous identifications are shown in red. The § symbol in the sample names shows when one species is indistinguishable from another and the * symbol is added when a species is absent in the deconvoluted possibility list. (**Bottom**) Overview of the experimental species content per sample. Details of the experimental design can be found in **Data S5**; the interactive output HTMLs are deposited in PRIDE (PXD079630).

### Sequence correction and reconstruction

Both MALDI-TOF- and LC-MS/MS-based palaeoproteomics rely on predicted protein sequences, which often are not validated via proteomics. False predictions, ranging from single amino acid mutations to largely incomplete sequences, are therefore common. We attempt to ameliorate this issue by leveraging peptide-patching for database correction in three different steps.

#### Point mutation database detection

We demonstrate this here for *Panthera pardus* (leopard), an important yet elusive representative of the genus *Panthera* during the Pleistocene in Europe. This species, according to its predicted sequence, carries two mutational sites. However, during validation using six individual *P. pardus* reference samples (Museum Koenig Bonn, Germany), we found that these mutations cannot be recovered, while we did recover the complementary peptides matching other *Panthera* species for all six individuals. In other words, these two predicted mutations are seemingly incorrect and a species-specific outcome for *P. pardus* based on the targeted COL1α1 and COL1α2 sequences is not possible, as shown by the presence of the complementary peptides. Therefore, all Panthera species cluster in this taxon in **Figure 3**.

#### Point mutation database correction

When an ‘X’ is present or built into a (higher taxonomic) collagen sequence, the correct amino acid can be filled out directly using the Mascot search engine, which considers all 20 amino acids when an X is present. For example, an X at residue 556 (COL1a1 sequence, *Loxodonta* genus) was identified as Glycine (G) based on a measured peptide match (data seen in **Figure S3**). While this approach needs additional validation, this particular mutation does match the peptide described by Buckley *et al*. (*26*).

Next, we noticed in several samples a persistently absent peptide that was otherwise consistently covered in closely related species and hypothesized that it could be indicative of a point mutation not present in the CollagenDB. In such cases, every amino acid of that peptide can be replaced by an ‘X’ consecutively in a single FASTA, thus allowing search engines like Mascot to fill in the blank residue. We verified this by examining the missing detectable peptide regions of the *Rhinoceros unicornis* (Indian rhinoceros) sample from the reference collection of Museum Koenig Bonn. This returned a Leucine at site 510 of COL1α2, producing a peptide not matching any closely related species in our database. Together with the mutational sites reconstructed within our pipeline, these mutations either match or extend on the *R. unicornis* sequences previously found by Welker *et al*. (*27*).

#### Extended Sequence Database correction

Using reference samples of known species, our approach can equally reconstruct longer missing and wrongly annotated sequence stretches. We demonstrate this on *Crocuta crocuta* (spotted hyena) reference samples: in the CollagenDB, the first 264 amino acid residues are missing in compare to *Hyeana hyaena* (striped hyena), and a wrongly predicted site is present in the middle of the sequence of COL1α2. First, the N-terminal sequence gap was peptide-patched with 102 measured amino acid residues from *H. hyaena*. Next, the wrongly predicted sequence ‘**FPEELMFFFSNSQ**GPAGVVGPVGAVGPR’ could be replaced with the measured peptide ‘**GEP**GPAGVVGPVGAVGPR’ from *Suricata suricatta* (meerkat). Similarly, in COL1α1 another wrongly predicted peptide was observed, namely ‘GDAGPPGPAGPTGPPG**STWLHALPQ**GNVGAPGPK’ which more likely should have been the measured peptide ‘GDAGPPGPAGPTGPPG**PI**GNVGAPGPK’ from *H. hyaena*. This specific measured peptide (and many other incorrectly predicted sites of *C. crocuta* COL1α1 and COL1α2, not described in detail, (**Data S3**)) was confirmed and is conserved in all samples within the *Hyaenidae* family.

### *Palaeoloxodon sp.* detection in mixed bags from Scladina Cave

Next, we analyzed dust from 24 bags storing >100 small bone fragments each, excavated in the locality of the Neanderthal child from Scladina Cave (complex of units 4A, which contains reworked Eemian (MIS 5e) sediments and material) (*12*, *28*, *29*). **Figure 5** shows an approximation of the potential candidates in each of the bags. We identified fauna matching the cave context, such as *Ursus sp.* (bear) and *Canis lupus* (wolf), as well as *Cervus sp.* (deer), *Equus caballus* (horse), and interglacial species such as *Sus scrofa* (wild boar)(*30*). Other species within Chiroptera, Rodentia and Eulipotyphla perhaps indicate that residual bone dust and soil contribute to the final species list, which in time could become an application in its own right: sampling cave soil for reconstructing the palaeofaunal spectrum of a given time period (*31*). Irrespectively, while the algorithm can approximate the content of mixtures quite reliably, it remains important to always balance the number of species considered and the reliability of the lower-scoring annotations. Highly complex mixtures from large mixed bags will always require careful inspection and individual verification of the candidate species.

**Figure 5:**
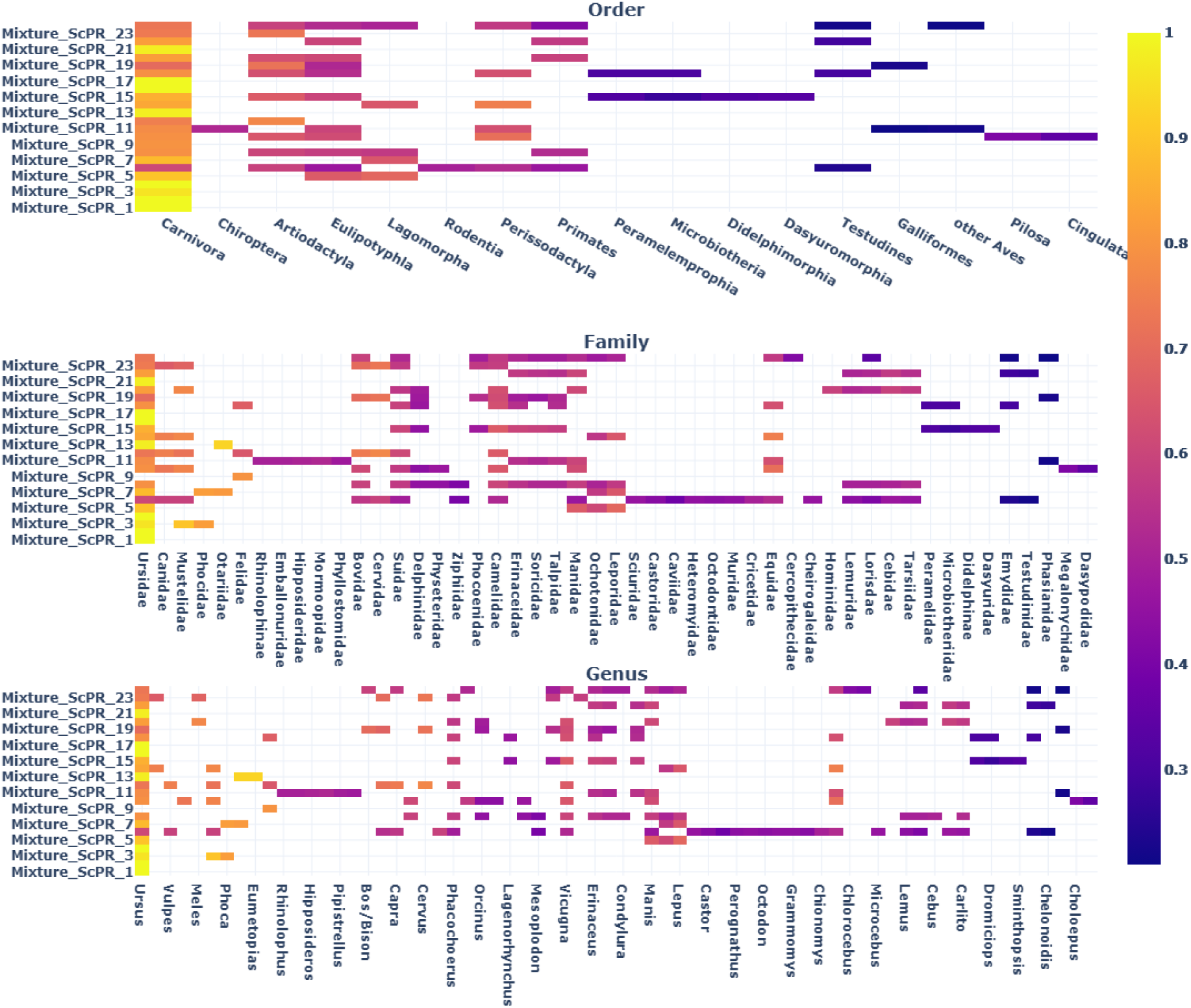
Analysis of bone dust from bags containing many remains reveals an overview of species present at Scladina. Bag dust was collected from bags related to the MIS 5e. The plot shows the approximation of potential species in each of the bags ‘Mixture_ScPR_’. Subplots show order-, family-, and genus-level approximations. This provides a bird’s eye view of the content of the bags (often >100 bone fragments per bag) and therefore of the palaeofauna of the Eemian in Scladina Cave in Belgium. The interactive HTML is provided as **Data S4**.

This illustrates not only the potential of the pipeline to investigate palaeodiversity via bulk analyses, but also helps to target specific species of interest. As a first example, some bag dust samples showed a very low score for bird remains. To assess the validity of this signal, several bone fragments were selected and separately sampled: one of the identifications was *Anserinae* (swan/goose; Sc-86-62-9). In a second example, three bags showed a positive signal for the family *Hominidae*, which could indicate the presence of fragments belonging to *Homo neanderthalensis* (Neanderthal). Therefore, these two bags were re-inspected and several individual bone fragments (*n=*32) from the bags were drilled and analysed. These almost exclusively turned out to be *Ursus sp.*, as this cave functioned as a cave bear den. Apart from two bone fragments belonging to *Cervinae*, we found a single bone fragment (Sc-86-82-2) that originated from a *Palaeoloxodon antiquus* (straight-tusked elephant), for which the recovered sequences match the *Palaeoloxodon sp.* sequences from Buckley *et al*. (*26*). This flat bone fragment could not be taxonomically assigned based on osteomorphological characteristics, however, retrospective histological features of the damaged cortical surface suggest that the specimen may derive from a juvenile. This also highlights the limits of complex bone dust mixtures, as no *Palaeoloxodon sp.* signal was captured in the dust of this particular bag, which is at least in part attributable to the overwhelming number of *Ursus sp.* fragments present. Still, another bag from the same locality did show low signals of *Palaeoloxodon sp.*, hinting that more bone fragments linked to this species could be present in this large assemblage. Although not unique within the broader geographic and chronostratigraphic context, this still represents the first and currently only specimen assigned to *Palaeoloxodon sp*. from Scladina Cave, as no other Elephantidae remains from the site have previously been attributed to this genus based on osteomorphological characteristics.

## Discussion

Here, we introduce the first tool for bulk proteomics analysis of multiple-species samples, using a novel ‘peptide patching’ method for sequence reconstructions. We hereby vastly extend on the application of collagen measurements in palaeoproteomics using the ClassiCOL algorithm. By approaching every sample as a patchwork of peptides derived from different closely related species, both physical and genetic mixtures can be deconvoluted. Relying on phylogenetic relatedness and slow evolutionary mutation rate, we were able to classify numerous and varied species and reconstruct missing collagen protein sequences in the database based on measured data. This allowed the reconstruction of palaeobiodiversity via bulk protein analysis of over 2,500 bone fragments in a single batch run, discovering the first straight-tusked elephant (*Palaeoloxodon antiquus*) found at Scladina Cave in this process, dating back to the Eemian (MIS 5e). The novel collagen sequences composed in this research can be used in future studies and classification efforts, which will present opportunities to expand collagen databases with additional taxa. Beyond future palaeontological research, we anticipate its application to other fields, including wildlife forensic studies.

## Methods

### Sterre collection (Ghent University, Belgium) samples

Carnivore samples from the reference collection of Ghent University included: *Phoca vitulina* bone (*n*=1), *Pusa sibirica* dentine (*n*=1), *Crocuta crocuta spelaea* dentine (*n*=2), *Panthera leo* bone (*n*=1) and *Hyaena hyaena* bone (*n*=1).

Artiodactyla specimens from the same reference collection included: *Potamochoerus porcus* skull (*n*=1), *Phacochoerus aethiopicus* bone (*n*=1), *Gazella dorcas* bone (*n*=3), *Boselaphus tragocamelus* dentine (*n*=1), *Tragelaphus strepsiceros* bone (*n*=1), *Tragelaphus scriptus* bone (*n*=1), *Tragelaphus spekii* bone (*n*=3), *Kobus kob thomasi* bone (*n*=1), *Hippotragus niger* bone (*n*=1), *Ammotragus lervia* bone (*n*=2), *Connochaetes gnou* bone (*n*=1), *Sylvicapra grimmia* bone (*n*=1), *Aepyceros melampus bone* (*n*=1), *Bison priscus* bone (*n*=1), *Giraffa camelopardalis* bone (*n*=2) and *Capreolus capreolus* antler (*n*=1).

Other specimens collected included: *Equus quagga burchellii* bone (*n*=1) and *Mammuthus primigenius* ivory (*n*=1). Detailed description and results can be found in **Data S5**.

### Ledeganck reference samples

Experimental physical mixtures were created by mixing drilled bone powder from ten different species from the Ghent University Ledeganck reference collection. The mammals selected were: *Rangifer tarandus, Sus scrofa, Capreolus capreolus, Ovis aries, Cervus elaphus, Vulpes vulpes, Meles meles, Martes foina, Lepus sp., and Mustela putorius*. 10 mg of cortical bone powder was drilled from each of the samples, taken from the left femoral midshaft. 14 unique mixtures were made by combining 1µl of the original peptide stock solution of each individual sample. A detailed description of the mixture compositions and results can be found in **Data S5** and **Figure 4**.

### Scladina Cave bag dust samples

Dust was collected from 24 individual bags containing varying numbers of bone fragments or tamisage (small fragments retrieved through wet sieving). All bags belong to the complex of units 4A of Scladina Cave. Detailed description and results can be found in **Data S5**.

### Samples from LIB-Museum Koenig Bonn (ZMFK), Germany

The bone and ivory samples from the collection of Museum Koenig Bonn (Germany) were donated by research institutes and zoos. This included *Panthera tigris sumatrae* (*n*=1) and *Panthera pardus ciscausica* (*n*=1) from the Wilhelma Zoo in Stuttgart. Also *Panthera tigris sumatrae* (*n*=2), *Panthera uncia* (*n*=1), *Panthera pardus* (*n*=2) and *Panthera onca* (*n*=2) from the Wuppertal Zoo. The collection also contained *Panthera uncia* (n=2) and *Proteles cristata* (*n*=1) from the Krefeld Zoo and *Proteles cristata* (*n*=1) from the Zoo of Hamburg. *Panthera pardus* (*n*=1), *Panthera onca* (*n*=1), *Panthera saxicolor* (*n*=1) and *Hyaena brunnea* (*n*=1) came from the Zoo of Köln and a *Hyaena brunnea* (*n*=1) from the Zoo of West Berlin. In addition to specimens donated by Zoos, several species had a wild provenance: *Panthera saxicolor* (*n*=1, Iran), *Crocuta crocuta* (*n*=2, Serengeti National Park, Tanzania), *Crocuta crocuta* (*n*=1, Cameroon), *Proteles cristata* (*n*=1, South Africa), *Loxodonta cyclotis* (*n*=3, Cameroon), *Elephas maximus* (*n*=1, Thailand), *Elephas maximus* (*n*=1, Sri Lanka), *Loxodonta africana* (*n*=1, East Africa) and *Diceros bicornis* (*n*=1, East Africa). Other samples with unknown provenance are *Loxodonta africana* (*n*=3), *Rhinoceros unicornis* (*n*=1), *Dugong dugong* (*n*=1) and *Ceratotherium simum cottoni* (*n*=1). The final sample from Museum Koenig Bonn collection was an *Elephas maximus* (*n*=1) specimen donated by the Anatomical Institute University Köln. The original museum sample numbers and additional sample metadata are listed in **Data S5**.

### Protein Extraction

An adaptation was made to the suspension trapping protein extraction protocol used in Engels *et al.* 2025 (*32*). From each sample, 5mg of bone was demineralized in 600 µL 0.6M HCl (ChemLab, CL05.0312.1000) overnight at RT, while shaking (750 rpm, Eppendorf Thermomixer comfort). This was followed by pelletizing the samples via centrifugation (10,000 rcf, 10 min, Eppendorf centrifuge 5417R), after which the pellet and supernatant fractions were separated. The pellet fraction was washed with ice-cold acetone (Sigma-Aldrich, 179124–1L). Trichloroacetic acid (TCA) was added to a final 1:3 v:v to the supernatant fraction and incubated on ice for 1h. After centrifugation, the TCA-HCL supernatant was discarded and the pellet washed with ice-cold acetone. Both pellets were resuspended in extraction buffer (5% SDS (Invitrogen, 15553–027) + 50 mM TEAB (Sigma-Aldrich, 90360–100 ML)) and combined. Disulfide bridges were reduced by addition of DTT (Chemlab, CL00.0481.0025) to a final concentration of 0.02M and incubated in the dark at 37°C for 30min. Proteins were precipitated with phosphoric acid (Chemlab, CL00.0605.1000) at pH 1.

HiPure Viral Mini columns (Magen Biotechnology, China; C13112) were used for the protein trapping. Binding/washing buffer (165µL, 100 mM TEAB in 90% Methanol (Chemlab, CL00.1377.1000)) was added to the sample before loading on the column. Reloading of the sample was performed to reduce degraded protein loss. This was followed by three washing steps by adding 150µL washing/binding buffer and subsequent centrifugation steps (4000 rcf). An on-filter protein digestion was performed using 0.5µg/40µL trypsin/Lys-C (ProMega, V5073) in 50mM TEAB for 2h at 47°C.

Peptide elution was performed by sequentially adding of 30µL of 50mM TEAB, 30µL of 0.1% formic acid (FA) (Biosolve, 2324), and 30µL of 50% acetonitrile (Chemlab, CL00.0194.1000) separated by 1 minute of incubation time and centrifugation (4000 rcf, 23°C). The samples were vacuum-dried for storage at −20°C before resuspension in 0.1% FA for LC-MS/MS analysis.

### LC-MS/MS Data Acquisition

LC-MS/MS data were acquired using the Waters Acquity M-Class UPLC system coupled with a Sciex ZenoTOF 7600 mass spectrometer using data-dependent acquisition mode. Peptides were trapped using a YMC triart C18 guard column, 3 μm, 5 × 0.3 mm and as an analytical column a YMC Triart C18, 3 μm, 150 × 0.3 mm was used for separation, using a non-linear gradient (20min, 1.5-to-36% solvent B (0.1% FA in acetonitrile) in solvent A (0.1% FA in water)). Precursor scans (MS1) over a mass range of 300–1600 m/z were acquired for 0.1s. A selection of up to 40 precursors, with an intensity threshold of 150 counts per second, and a dynamic exclusion of 6s after 2 occurrences. Collision induced dissociation (CID) was used to fragment precursors with charge states between 2 and 5 per cycle. Fragment spectra acquisition (MS2) was done for 0.015s over 100-2000 m/z mass range, with a cycle time of 0.920s.

### Peptide identification

Peak-picking was performed by use of MSConvert, converting the raw *.wiff file into a MGF file format. Peptide-to-spectrum matches were identified using Mascot Daemon (version 3.0.0, Matrix Science, London, U.K.), and searched against the CollagenDB (reference samples were restricted to Mammals) from Engels *et al*. 2025 (*11*), and a Universal Contaminants database (*33*). Enzyme parameter was set to semiTrypin and 1 missed cleavage, and variable modifications Deamidation (NQ, + 0.984016 Da) and Oxidation (MP, + 15.994915 Da) were used as search parameters alongside a fragment error tolerance of 30 ppm and peptide tolerance of 10 ppm.

### ClassiCOL and the CollagenDB

As input to the isoBLAST-ClassiCOL pipeline (*11*), a Mascot CSV result file was generated setting a significance threshold of p < 0.01. The ClassiCOL search was restricted to Mammalia for individual reference samples (Sterre and Bonn). All taxa in the database were considered while searching the mixtures. Default settings were used for the classification.

### Collagen phylogenetic tree construction

Protein sequences of COL1α1 and COL1α2 were extracted for all species in the CollagenDB under the Laurasiatheria. Species with a lower than 85% average sequence length were discarded from the analysis, resulting in 113 species. This was followed by concatenation of the collagen protein sequences after which a multiple sequence alignment was performed using the multiple alignment part of the ClassiCOL code described below. This multiple alignment was exported in FASTA format and loaded into the MEGA11 software (*23*). In this step ambiguous regions, sequence stretches covering >90% of the species with gaps and prediction mistakes (peptides <5% similarity compared to the whole species set) were removed. Muscle re-alignment was performed and a phylogenetic tree was inferred using the Neighbour-Joining method (*34*), and a bootstrap consensus tree was built (*n*=500) (*35*). A Poisson correction method was used to infer the evolutionary distances (*36*).

The multiple sequence alignment can be used to define the uniqueness of detected amino acid residues for extant relatives present in the database and therefore the probable ancestral origin of the sample. Each of the detected potential mutational residues are therefore scored for *recentness* of the mutation according to their ancestral lineage and uniqueness compared to other candidates in a one-vs-one comparative approach.

As expected, the number of mutational residues compared to one given species gradually goes up towards the last common vertebrate ancestor (LCVAs). This is the *in-silico* equivalent of going further back in time. With the substantial differences in generational turnover between e.g. Artiodactyla and Rodentia, their collagen mutation rates therefore also vary considerably (**Figure S2**). By normalizing these rates separately however, this approach co-ordinately accounts for less obvious processes that lead to altered evolutionary pressure on the protein.

Building on this evolutionary logic, we attribute a gap in collagen sequence coverage to the absence of the target species in the database and not specifically to our inability to detect ‘gap’ peptides via LC-MS/MS. Indeed, if a given peptide is usually detected in most species, its absence in a particular sample is more likely due to absence from the database. Building on this, residues of closely related species can be used to ‘patch’ these gaps to create a theoretical sequence, i.e., a sequential combination of existing but not co-occurring collagen peptides, which approximates the true species more closely. By iteratively comparing each new theoretical sequence with the remaining sequences from the previous iteration, the algorithm moves upstream in the taxonomic tree towards the LCVA, in a one-vs-one approach, which shifts into a many-vs-many approach near convergence. Irrelevant theoretical combinations can be filtered out because of their lower **combinatorial summarized residue score** (**Formula S1**) compared to a the LCA and because deterministic peptides can be found in other lineages when approaching the LCA. In other words, the data pushes the score towards the LCA rather than to a missing species under the LCA. This convergence ends with either an exact match to a species in the database or with a theoretical sequence which can be localized to a phylogenetic location adjacent to close relatives, providing a **theoretical candidate**. When dealing with a genetic mixture, this convergence reflects the complementarity of missing residues in a close relative that can be filled by another relative.

When dealing with true physical mixtures, i.e. a single sample containing multiple species, multiple explanations are found in the data for a given location in the collagen LCA consensus sequence. Thus, there are different peptides that cover the same sequence stretch. Still, as we isobarically ambiguate peptide sequences during the isoBLAST run, overlap can also occur due to data ambiguity. Therefore, both theoretical candidates and potential physically mixed candidates are considered for the next iteration until convergence.

Finally, the most challenging case is a physical mixture containing genetic mixtures (e.g. a multiple-species sample in which at least one taxon is extinct), which could lead to an even higher number of multiple output candidates, like the physical mixture, but with an increased number of theoretical candidates.

In each case, a distance matrix can be calculated on the measured data only, containing the distances between all species in the database and the newly formed theoretical candidates. For this, each measured distance can be extrapolated onto the taxonomic distance matrix of the NCBI taxonomic tree, resulting in a matrix containing the measured relationship amongst the potential candidates and their closest relatives. The numerical distances can then be used to calculate the distance between the theoretical candidates and taxa linked to the lineages of closely related species.

As an output, ClassiCOL_v2 provides two different perspectives (**Figure 2A and B**). The first is the clustering of different matches and theoretical sequences based solely on measured peptides. This provides an intuitive summary of the potential taxonomic regions that are present in the sample. Next, the algorithm generates a separate output from the perspective of each of these outcomes (theoretical and matching). This **taxonomic Slope Plot** plots the distances from the outcome to the rest of the tree. A steeper slope implies that a smaller taxonomic distance is covered towards the next taxon.

Taxonomic slope plots are generated from the same sample. In each header, the BC score ranks the candidates. Note that it makes little sense to preset a threshold for selecting potential species, because even low scores could still provide a lead in some samples where only a fraction of certain animals is present, e.g. in the dust from bags containing large collections of bones, thus a mixture. Yet it is reasonable to try to pinpoint score drop-offs when e.g. a single bone was drilled (a single genetic mixture for example). Rather, we advocate to turn to all the output files to make an informed decision.

### Mixture algorithm

#### Input data and defining the mutational search space

The mixture analysis is performed post-ClassiCOL analysis, starting from the potential species candidate list along with peptides recovered via isoBLAST. Each of the potential species candidates are grouped by their Order taxon. Then, up to three representatives are chosen for each of the taxa linked to the remaining candidates, and added to the Order list as positive control for potential missing mutations. Next, all sequences are extracted from the database that are linked to the peptide list, with a minimal amount of 25% protein coverage (e.g. if COL1a1 has a >25% coverage, all COL1a1 sequences will be taken to the next step). These sequences will be subjected to a quality check, which includes sequence length match (>90% minimal length, truncated sequences are discarded) and 25% of total peptide match (removes incorrect sequences). This is followed by a multiple sequence alignment per sequence group, using at least 2 species. When a single species under the Order level is in the output it will be separated as a single match. This multiple sequence alignment is performed using the taxonomic tree, where a 1vs1 sequence alignment is performed on each taxonomic level, starting from the genus level. Anchor points, being tryptic peptides, are used to better align these sequences. The resulting alignment is convoluted into a consensus sequence, replacing mismatches with an ‘X’. After iterative alignments up to the Order level, we return the mutational search space for each of the ‘Orders’ given their considered candidates. The search space is plotted and saved for easy manual curation of the search space by the user when needed. The resulting Order level multiple alignments are used for the following mixture analysis. This analysis starts with the mapping of the measured peptides onto the multiple sequence alignment, irrespective from which the species they originated.

#### Species inference the initial iteration

Before starting the mixture analysis, all orders are checked for uniqueness, removing those that do not show uniqueness at the order level. This means that the Order groups must have >10% unique peptides compared to the maximal amount in the data, and a minimum of 1.5 unique peptides per candidate at the Order level are required. The mixture analysis is an iterative process that is performed on each of the Order groups separately. For each of the groups, the initial iteration consists of grouping close relatives for each of the protein sequence types, in a 11 comparison. A backbone consensus sequence is taken from the multiple alignment matching the ancestral taxon of each comparison. This is followed by looking for complementarity of each of the mutational amino acid residues at the taxon LCA level. To achieve this, the algorithm iterates through all mutational residues in the consensus backbone and checks which residues are present in the measured data. Each of the mapped residues gets a unique score per iteration, which is the sum of the uniqueness of the mutation under the LCA and the chance it is a recent mutation. The uniqueness of the mutation is determined by the number of species in the database that have the measured residue divided by the number of species in the database and is adjusted for missingness of species in the database which could potentially contain the residue under the LCA. The recentness of the residue is determined by the number of possible amino acids at the residue location divided by the number of taxa under the LCA in the database species adjusted with the amount of missingness under the LCA. Each residue is assigned to the taxon where the first instance of the measured residue was noticed under the LCA.

After the 1VS1 comparisons are performed, a taxon contribution score is calculated to determine the chance that a species is present in the sample. This score is calculated for each of the comparisons as follows: all protein groups are concatenated and the measured mutational residues are filtered out globally. During the initial step, the potential for the comparison to consist of a mixture is considered, based on overlap in mutational measured residues. In other words, if different amino acids where measured for a single residue, the algorithm flags the taxa as potentially mixed. The uniqueness of the residue is normalized by their presence in other comparisons. After mixture flagging, a normalized summed residue score is assigned to each taxon in both lineages, linking the two species under consideration.

#### An iterative step for determination of physical -or genetic mixtures

Each of the comparisons can result in three possible end results. A) There is no difference between 2 candidates, or 1 candidate is a subset of the other. This will result in the removal of the subset (or one of the identical candidates) for the next iteration. B) A potential physical mixture is flagged. In this case the flagged candidate is retained as a potential final candidate. Additionally, when there is a possibility for a genetic mixture (i.e. complementary residues can be combined to a new sequence) this chimeric species will be assigned to a theoretical taxon using the common ancestor of the donors as starting point during point C. C) A genetic mixture is possible, meaning both candidates contain measured residues which are missing in the other. These candidates will be combined to a novel theoretical taxon for the next iteration. During each iteration theoretical candidates will be assigned a taxonomic lineage matching the calculated LCA. The 1VS1 comparisons will, similarly to the first iteration, be based on taxonomic proximity. Per candidate a single comparison with another candidate with an Order level LCA is allowed. In other words, if a candidate survives to this point it is most likely to be present in the sample.

#### Candidate proximity to a taxonomic lineage

The sequences of all remaining candidates, both database matches and theoretical candidates, are concatenated. Candidates that remain a subset of another or have an Order uniqueness lower than 2 peptides are discarded. A distance is calculated on the concatenated sequences between all candidates and species in the database under the Order taxa. A distance of 1 is given to all comparisons beyond the Order level. Next a distance matrix is calculated based on the known taxonomy, according to NCBI, providing a step wise distance between species (e.g. two species under the same genus will be given a step distance of 2). The theoretical candidates are extrapolated onto the step-distance matrix from the measured data by linking their distances to each taxonomic level in the closest related species lineage to the measured data distances. Additionally, missing taxa from the database can be inferred by calculating the theoretical distance from the missing LCA to the measured data and extrapolating it into the stepwise distances. These results are plotted in a tree-plot showing the measured distance relationships of the species in the database together with candidates. In addition, a line plot for each of the candidates is created which shows the extrapolated stepwise distances towards the considered and missing taxa.

#### Creation of FASTAs containing sequences based on measured data

All remaining candidate sequences are output in FASTA format in three distinct ways for easy handling by the user. 1) All measured peptides are mapped on the consensus sequences. Residues that were not measured will be returned as ‘X’. 2) All mapped peptides and residues that show no signs of mutation throughout the lineage are returned, an ‘X’ is assigned to unmeasured mutational residues. 3) the sequence of the scaffold protein in the database.

## Supporting information

DataS1_interactive_figure2

Data_S2_interactive_figure3

Data_S3_hyaenidae_multiple_alignment_measured_data

Data_S4_interactive_html_figure5

Data_S5_sample_metadata_and_overview

## Data availability statement

Raw data (Sciex WIFF), peak and search data (MGF, mzIdentML) and result files (CSV, ClassicOL_v2 output folders) have been deposited on the PRIDE(*37*) repository (ProteomeXchange, https://www.ebi.ac.uk/pride/) with identifier PXD079630. The ClassiCOL_v2 code and guidelines have been made available on GitHub (https://github.com/EngelsI/ClassiCOL). All supplementary data and figures can be retrieved in the supporting information folders and in the supplementary data file.

## Acknowledgements

The authors acknowledge the ProGenTomics Core Facility (www.progentomics.be) for performing the sample measurements and for providing expert assistance throughout the study. We would like to give a special thanks to the UGent Computational Biology class of 2024-25 for optimising the ClassiCOL_v1 code with the intent to minimise computational time.

## Author contribution

I.E, M.D., S.D. Conceptualisation; I.E., A.B., S.M.S. performed sample collection of Sterre material; I.E., A.B., G.A., M.D. performed sample collection at Scladina Cave; I.E., A.B., T.D. performed sample collection of Ledeganck reference collection; I.E., A.B. experimental design of mixture benchmark; I.E., S.D., A.T., J.D. performed sample collection of Museum Koenig Bonn material. J.D. reference collection supervision Museum Koenig Bonn; T.VdB. reference collection supervision Sterre, K.D.M. Scladina Cave sample supervision. I.E. coding and visualisation; S.VdV. computational time improvements ClassiCOL_v1; A.B. performed computational time comparison ClassiCOL_v1 vs ClassiCOL_v2; I.E. performed protein extraction; I.E., S.D. performed LC-MS/MS analysis; I.E. data analysis; M.D., D.D. project supervision, I.E., M.D. writing-original draft; T.D. writing-Palaeontological contribution. All co-authors reviewed and provided feedback on the manuscript.

## Data, code, and materials availability

The code and user guidelines have been made available on GitHub: https://github.com/EngelsI/ClassiCOL. Proteomics data files have been made available on ProteomeXchange/PRIDE with identifier PXD079630.

## List of Supplementary Materials

**Supplementary Materials (PDF):** contains supplementary figures and formulae described in the main text. **Figures. S1 to S3 and Formula S1**

**Data S1 (ZIP):** Interactive HTML files of figure 2

**Data S2 (ZIP):** Interactive HTML files of figure 3

**Data S3 (ZIP):** MEGA11 alignment files of Hyaenidae measured sequences

**Data S4 (ZIP):** Interactive HTML files of figure 5

**Data S5 (ZIP):** Excel document containing the results and sample metadata of all performed analyses.

