## Supplementary materials for "Palaeoproteomic deconvolution of physical and genetic collagen mixtures"

**Funding:** This research was funded by a grant from the Special Research Fund (BOF) of Ghent University with funding code BOF.GOA.2022.0002.003

**Keywords:** Palaeoproteomics, Bioinformatics, Palaeontology, collagen, Bioarchaeology

### Supplementary figures

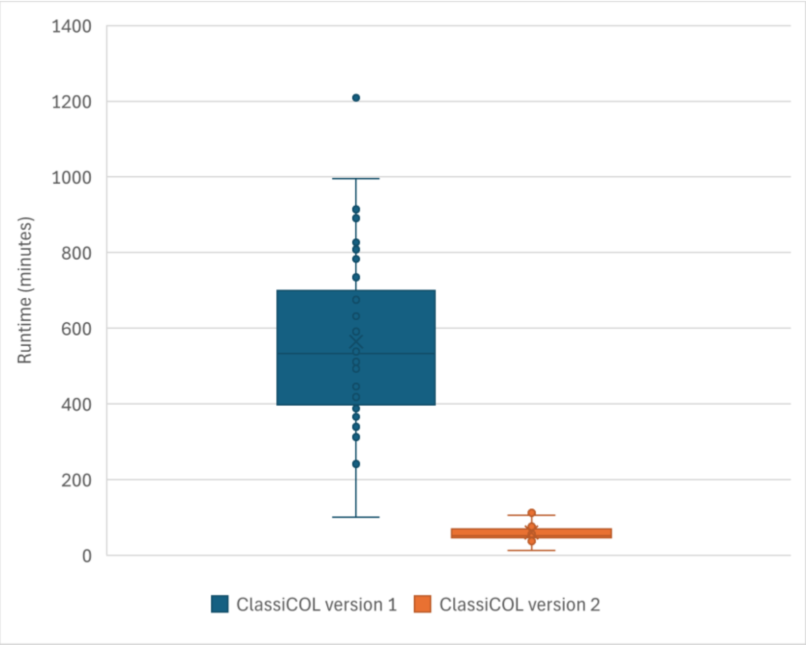


**Figure S1: Computational time requirement of ClassiCOL2 compared to the original version.** 42 samples with abundant collagen peptide content were analysed with the original ClassiCOL pipeline and again with the ClassiCOL2 pipeline. Time from submission to final file generation are plotted in minutes. All samples were analysed in a single batch submission. The plot shows an average 10 time reduction in computational time when using the newest version.

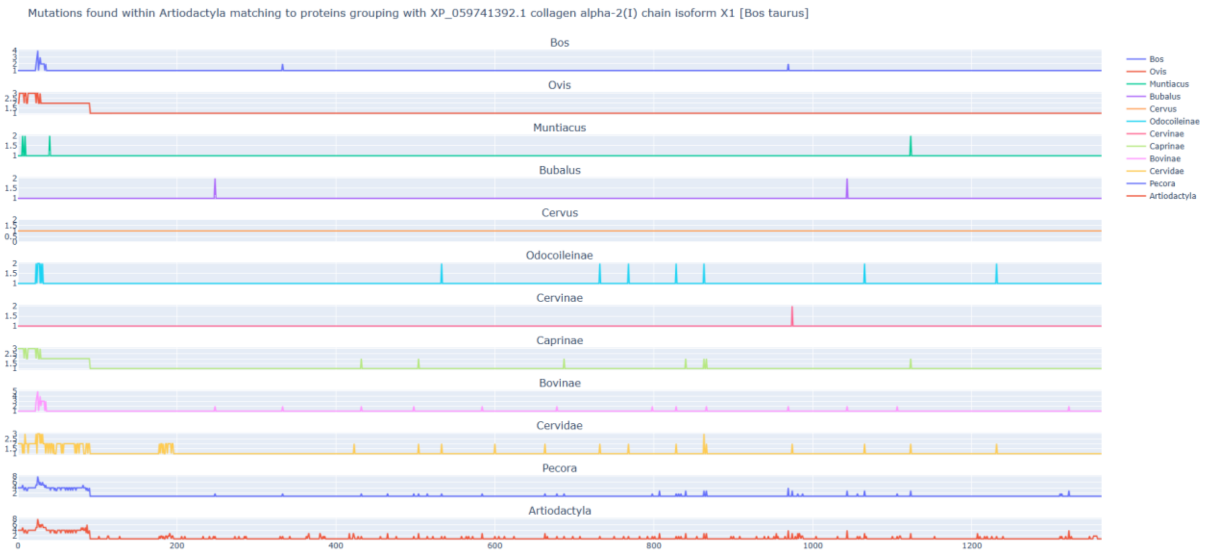

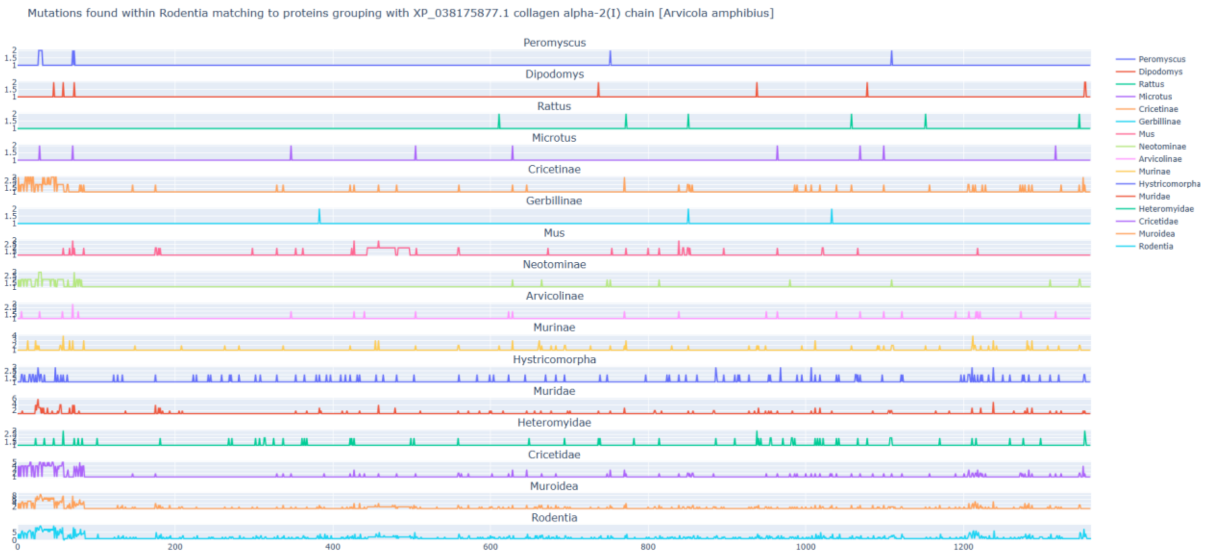


**Figure S2: There is an increase in mutational space when the generational turnover increases.** Plots shows the mutational space for the Laurasiatheria (top) and Rodentia taxonomic level (bottom). An increase in mutational residue location is visible comparing Rodentia to the Laurasiatheria.

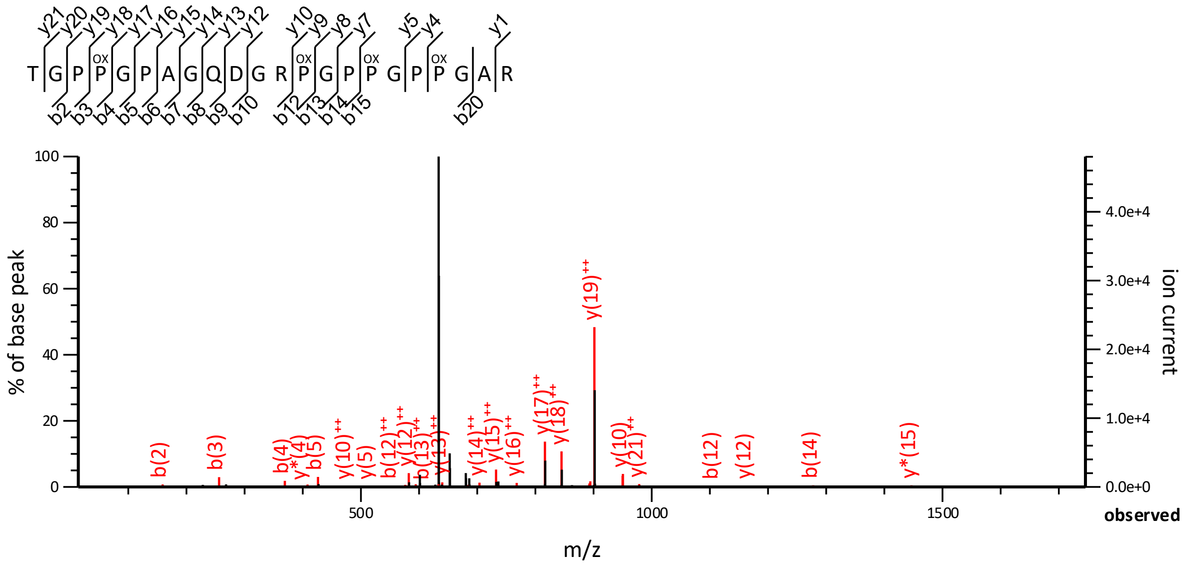


**Figure S3: Peptide spectrum match of *Loxodonta africana* sample with filled out 'X'.** The peptide spectrum match was found in the ED5 sample by use of Mascot. The original peptide as found in its protein sequence is TGPPGPAXQDGRPGPPGPPGAR. This spectrum thus shows the capability of search engines to account for missing amino acid residues together with the ion series coverage of the residue y15 ion and b8 ion.

### Supplementary formulae

**Formula S1 depicts this summarized residue score.** The summarized residue score is based on the uniqueness of the residue in the database and the theoretical recentness of the mutation. In other words the overall recentness of the mutation in relation to the taxa under investigation and the uniqueness in relation to the taxa that are being compared. This means that we check the chance that the residue is unique at each taxonomic level towards the LCA of AB. Very similar to P_unique but with the difference that it sums the chance at each taxonomic node. The residue score is thus the discriminatory score of the residue when comparing taxa A and B. Taking into account that the unique mutation might be shared amongst others in the same lineage.

$$P_{recent}(A|B) = \sum_{T \to [AB, lca[} \left(1 - \frac{\sum species\ in\ DB\ under\ T\ with\ same\ aminoacid\ residue}{\sum species\ in\ database\ related\ to\ T}\right)) * (1 - (\frac{\sum species\ missing\ under\ T}{\sum all\ species\ known\ under\ T})$$

$$P_{unique}(A|B) = 1 - \left(\frac{\sum species\ in\ DB\ with\ same\ aminoacid\ residue}{\sum species\ in\ database\ related\ to\ LCA_{AB}}\right) + (1 - (\frac{\sum species\ missing\ under\ LCA_{AB}}{\sum all\ species\ known\ under\ LCA_{AB}})$$

$$Residue_{score} = P_{unique} + (\frac{P_{recent}}{distance\ to\ LCA})$$

$$Summarized\ residue\ score\ per\ taxon_{n \to LCA} = \frac{\sum residue\ score\ taxon_n}{\sum residue\ score\ ALL}$$
