## Supplementary figures and images for "Palaeoproteomic deconvolution of physical and genetic collagen mixtures"

### bag_dust_output_scladina.png

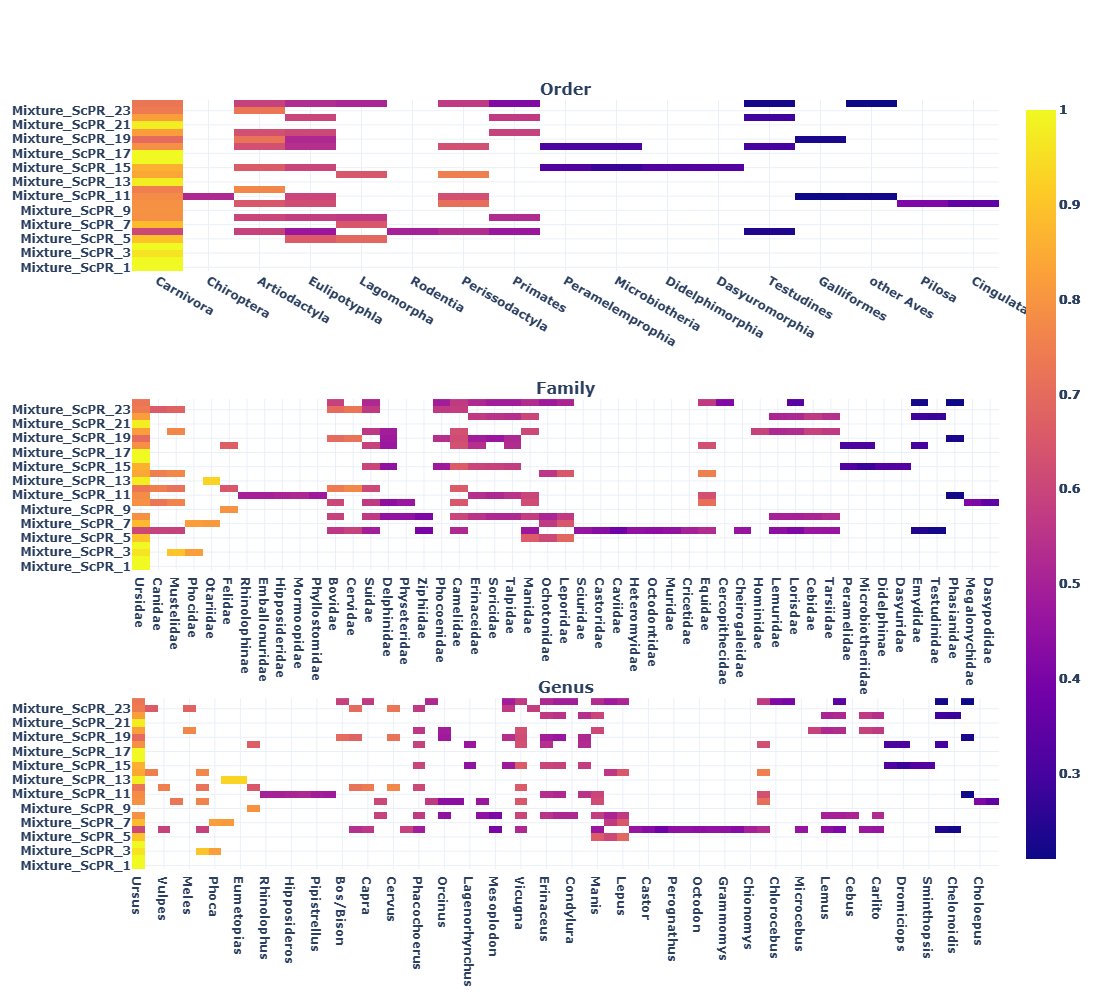

### benchmark_overview_plot_bonn_sterre.png

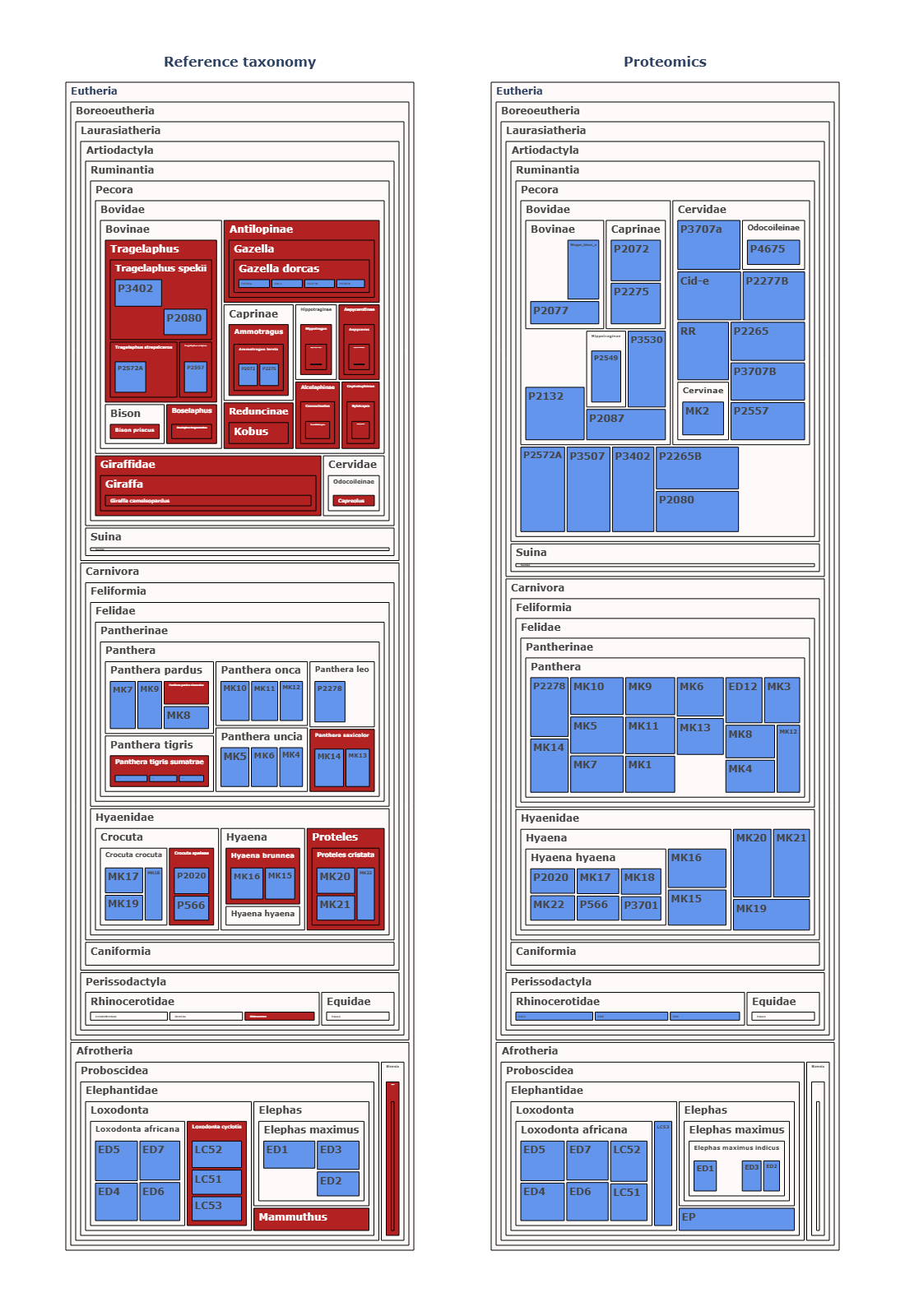
